# GRAPHENE OXIDE AS A NOVEL IMMUNOTHERAPY TOOL FOR THE MODULATION OF MYELOID-DERIVED SUPPRESSOR CELL ACTIVITY IN THE CONTEXT OF MULTIPLE SCLEROSIS

**DOI:** 10.1101/2023.03.28.534540

**Authors:** Celia Camacho-Toledano, Isabel Machín-Díaz, Rafael Lebrón-Galán, Ankor González-Mayorga, Francisco J. Palomares, María C. Serrano, Diego Clemente

**Author notes:** Current address: Hospital Universitario de Toledo, Avd. Río Guadiana, s/n 45004. Toledo, Spain. Current address: Hospital Universitario de Navarra, C/Irunlarrea 3, 31008 Pamplona, Navarra, Spain (Ankor González-Mayorga).

## Abstract

Multiple Sclerosis (MS) is a chronic, inflammatory disease of the central nervous system. Despite the pharmacological arsenal approved for MS, there are treatment-reluctant patients for whom cell therapy appears as the only therapeutic alternative. Myeloid-derived suppressor cells (MDSCs) are immature cells of the innate immune response able to immunosuppress T lymphocytes and to promote oligodendroglial differentiation in experimental autoimmune encephalomyelitis (EAE), a preclinical model for MS. Culture devices need to be designed so that MDSCs maintain a state of immaturity and immunosuppressive function similar to that exerted in the donor organism. Graphene oxide (GO) has been described as a biocompatible material with the capacity to biologically modulate different cell types, including immune cells. In the present work, we show how MDSCs isolated from immune organs of EAE mice maintain an immature phenotype and highly immunosuppressive activity on T lymphocytes after being cultured on 2D reduced GO films (rGO_200_) compared to those grown on glass. This activity is depleted when MDSCs are exposed to slightly rougher and more oxidized GO substrates (rGO_90)_. The greater reduction in cell size of cells exposed to rGO_90_ compared to rGO_200_ is associated with the activation of apoptosis processes. Taken together, the exposure of MDSCs to GO substrates with different redox state and roughness appears as a good strategy to control MDSC activity *in vitro*. This versatility of GO nanomaterials and the impact of their physico-chemical properties in immunomodulation open the door to its possible selective therapeutic use for pathologies where MDSCs need to be enhanced or inhibited.

---

Multiple sclerosis (MS) is a chronic neurological disease affecting 2.5 M people worldwide. In MS, the harmony of immune tolerance related to the nervous system is broken. As a consequence, the immune system attacks the brain and spinal cord producing demyelination and neurodegeneration, which leads to sensory, motor and cognitive symptoms and a gradual increase in patient disability^1^. 85% of MS patients experience the relapsing-remitting MS (RRMS) in which periods of symptoms exacerbation are followed by full or partial recoveries^2^. Due to an extensive investigation in MS therapy, over 18 disease modifying treatments (DMTs) are available for RRMS devoted to control relapses and slow the progression of disability^3^. However, in an important proportion of RRMS patients showing early and highly active MS, the risk of disability accumulation is high and for many of them current DMTs provide suboptimal efficacy, with cell therapy being the only alternative^4, 5^. In this context, regulatory cells of the innate and adaptive immune responses appear as a suitable therapeutic target to control the exacerbated immune response, favor the restoration of immune tolerance, and induce nervous tissue regeneration^6^.

In the last years, the study of myeloid-derived suppressor cells (MDSCs), a highly immature regulatory cell type of the innate immune response, is gaining importance in the context of MS^7, 8^. MDSCs have been shown as key elements in the control of symptoms recovery in the murine MS model, experimental autoimmune encephalomyelitis (EAE)^9^. In fact, MDSCs are being identified as a future target for new MS interventions due to their multifaceted roles as orchestrators of the control of the adaptive immune response (*i.e.,* T cells)^10–13^, together with their involvement in favoring myelin restoration^14^ and preventing the harmful microbiota disbiosis induced by EAE^7^. *In vivo* studies in EAE have shown that the maintenance of the number and immature state of MDSCs are crucial for them to exert their potent immunosuppressive action and control disease severity^13, 15^. However, once isolated from the donor organism, MDSCs tend to differentiate spontaneously, which greatly impairs their potential usefulness for future cell therapies that require their isolation and prior expansion on standard cell culture conditions. For this reason, research is needed on new materials for cell growth whose interaction with MDSCs keeps them immature and highly immunocompetent.

Nanomedicine has emerged as a revolutionary field to provide novel customizable therapies in the nanoscale for a wide plethora of medical applications. Specifically, it makes use of nanosized tools for the diagnosis, prevention and treatment of diseases^16^. The interest is such that an increasing number of applications and products containing nanomaterials, or at least with nano-based claims, have being made available to date. Within those, nanosized particle-based platforms for immune-related biomedical applications are a relatively novel field in which both immunosuppression and immunoactivation are being pursued, including immunotherapy tools, vaccine carriers, adjuvants, and drug delivery systems to target inflammatory cells. Among the most relevant nanomaterials under investigation for biomedical applications, graphene-derived materials (GDMs) are becoming promising candidates for both diagnostic and therapeutic uses. Regarding immune-related applications, the exploration of GDMs is limited and mainly focused on three major topics: (i) Their interaction with immune cells for systemic biocompatibility assessment and the induction of specific immune responses, (ii) The development of immune-biosensors and (iii) Their use in combination with antibodies for tumor targeting^17^. In the first case, although largely unexplored until recently, GDMs are displaying a surprisingly attractive ability to interact with immune system elements, either by stimulating or suppressing specific responses. To this regard, their different physico-chemical properties including purity, shape dimension, oxidation degree, and functionalization are essential drivers of this immune interaction. However, and despite discrete progress in the field, the role that these properties of GDMs play on their specific interaction with immune cells is still poorly understood.

Recent advances on the exploration of GDMs, including graphene oxide (GO), on immune cell interactions induce important immunosuppressive actions over different myeloid cells^18^, being more extensively explored in the case of macrophages^19^. Particularly, GO seems to have an outstanding ability to induce macrophage cytotoxicity, but also to alter their phagocytic capacity and, very importantly, polarize them towards either pro-or anti-inflammatory phenotypes by manipulating different physico-chemical properties of this nanomaterial such as its oxidation degree^20, 21^. Additionally, GDMs are able to modulate mesenchymal and neural stem cell differentiation *in vitro*^22^, thus bringing an alternative therapeutic potential to these nanosized carbon materials for their use in cell-based treatments as culture devices to either favor or hamper cell differentiation.

Herein, we describe the pioneer use of GO-based 2D culture substrates for the modulation of MDSC differentiation, which prevents the loss of their highly immunosuppressive activity over activated T cells in the context of the murine model of MS. Moreover, we identify that the oxidative degree and roughness of GO exerts a pivotal role on cell shape and differentiation, immunosuppressive activity, and viability.

## RESULTS AND DISCUSSION

### Fabrication and characterization of rGO substrates for MDSC culture

GO substrates for MDSC culture were fabricated by spin coating as thin layers on top of glass coverslips from a commercial GO slurry. GO sheets in the slurry were composed of a few layers of carbon, with lateral sizes at the micron scale as proved by transmission electron microscopy (TEM; **Figure 1A**). After thermal reduction at 200 °C, the resulting films (rGO_200_) were visualized by scanning electron microscopy (SEM) confirming that rGO sheets compacted and extensively covered the surface of the glass coverslips as a continuous coating with irregular wrinkles of diverse height and length (**Figure 1B**). The specific roughness of such rGO coatings were measured by Atomic Force Microscopy (AFM; **Figure 1C**). rGO_200_ substrates were characterized by *R_q_* values of 38.8 ± 4.26 nm, *R_a_* of 30.4 ± 3.37 nm and *R_max_* of 290.1 ± 38.54 nm. Next, a careful chemical characterization was performed. **Figure 1D** displays the comparison between X-ray Photoelectron Spectroscopy (XPS) spectra of the C 1s core level of the GO slurry and its rGO counterpart annealed at 200 °C (rGO_200_). Spectra were normalized to the maximum intensity to highlight line shape differences, which provides direct valuable insights regarding the chemical environment of C atoms. The elemental content of C and O for the different samples is provided in **Table 1**, as well as the relative percentage of the different C-O functional groups. From the quantitative analysis of XPS peaks upon sample anneals, it is observed that the O *versus* C atomic content ratio was reduced from 0.86 (GO slurry) to 0.10 (rGO_200_). This fact is also in line with the signal in the energy region of C 1s where the photoelectrons from C-OH and O-C-O groups are emitted, whose intensity also decreased with the O 1s signal (see below). When analyzed in detail, XPS spectra of GO exhibit its characteristic photoelectron emission comprised of two wide and intense peaks in the binding energy range from 282 to 290 eV. Detailed peak shape analysis was performed by the deconvolution of the C 1s spectrum with several Gaussian/Lorentzian symmetric components (ratio of 85/15) using a least-squares fitting routine. The energy position of the peaks and their relative heights were determined to account for the emission ascribed to the different chemical environment of carbon atoms according to the values reported in previous work^23, 24^. The result of this fit provides a single symmetric peak from C-C emission representative of the oxide nature of GO and several components attributed to the presence of various oxygen functional groups denoted as Σ(C-O) in **Table 1**. On the contrary, C 1s emission from sample rGO at 200 C displays a very different lineshape dominated by an intense and asymmetric peak centered close to 284.5 eV. In addition, the long tail on the high energy side of the main peak usually reveals the metallic character of the sample, which certainly supports its graphitic-like nature and ascribes to sp^2^ bonding. Then, core level fitting of the spectrum is done by the deconvolution of an asymmetric component characteristic of C sp^2^ bonding (whose parameter values-peak position and full width half maximum-were obtained by fitting the C 1s signal from a HOPG reference sample) besides the symmetric components from the functional groups Σ(C-O). In this case, curve fitting of the whole spectrum makes it hard to clearly identify the sole emission from C-O and C=O chemical states, so their full contribution is added in **Table 1**. Moreover, a weak component appears, broader that any of the previous ones,, shifted at a higher binding energy value of ca. 6.5 eV from the main C 1s peak, which is clearly linked to the characteristic π-π* shake-up transition consistent with the majority existence of sp^2^ bonding in the sample. Note that a small amount of disordered regions where C atoms with defect-like sp^2^/sp^3^ bonding nature might also exist and those are included in the C-C component, but with no significant weight due to the low chemical shift and their minor contribution.

**Figure 1.**
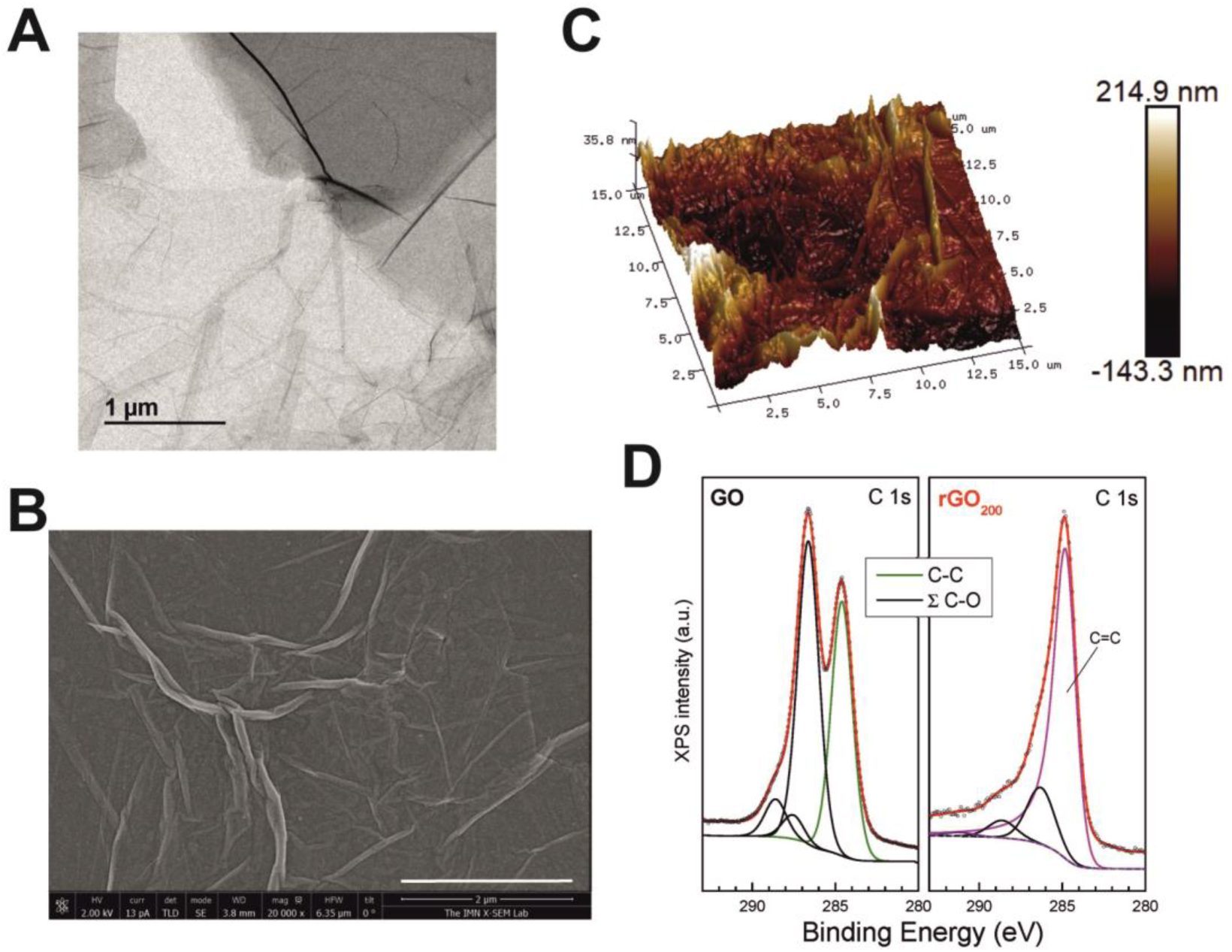
Physico-chemical characterization of rGO_200_ substrates for the modulation of MDSCs phenotype in vitro. (A) Representative TEM image of GO nanosheets in the slurry. (B) Representative SEM images illustrating major topographical surface features of the film. (C) 3D surface plots as measured by AFM. (D) XPS spectra of the C 1s corresponding to GO slurry (left) and rGO_200_ (right) together with their corresponding fits composed of chemically shifted components associated to different C chemical bondings. XPS spectra were normalized to the maximum peak intensity in each case for a better visual inspection and direct comparison among samples in order to highlight line-shape differences and contribution for each C chemical environment. Data points in every spectrum are represented as open symbols and Shirley background and component peaks using solid lines. The fitting curve (red line) resulted from the addition of several contributions belonging to: C-C bonding for GO (C-H, C-C defective sp_3_/sp_2_ configurations; green), graphitic structures for rGO_200_ (magenta), oxygen functional groups grouped as ⅀(C-O) (C-OH carbonyl, O-C-O carbonyl and O-C=O carboxylic groups at the energy shifted values reported in the literature;black) and p-p* transitions coming from graphitic structures for rGO_200_ at binding energy of ca. 291.0 eV (magenta).

**Figure 2.**
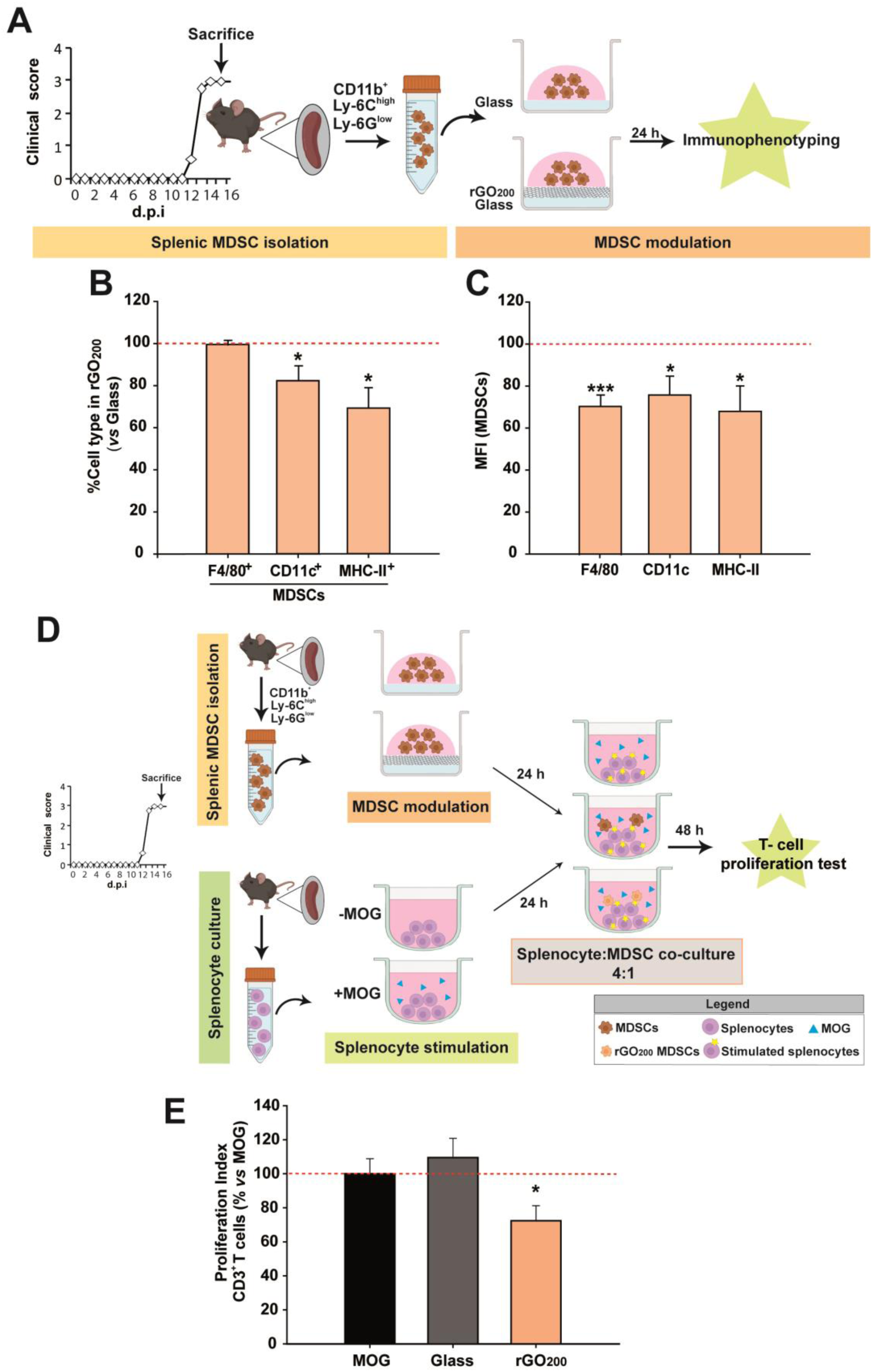
The immunosuppressive capacity of SP-MDSCs is enhanced after culture on rGO_200_ substrates by preserving their undifferentiated state. (A) Schematic representation of the experimental procedure to test rGO_200_ effect on MDSC immunophenotye. (B-C) rGO_200_ blocked the cell percentage (B) and MFI (C) of SP-MDSCs maturation markers compared to glass coverslips (red dashed line). (D) Schematic of the experimental procedure to test the effect of rGO_200_ on MDSC immunosuppressive activity over MOG-stimulated splenocytes from EAE mice. (E) Immunosuppressive activity of SP-MDSCs over encephalitogenic T cells after culture on rGO_200_ (red dashed line represents the proliferation observed after MOG stimulation). Data are shown as the mean ± standard error of the mean of N=3 in all experiments.

**Table 1.** O/C atomic ratios determined from O 1s and C 1s core levels emission. Relative percentage of peak areas for each type of C bonding obtained by XPS C 1s analysis.

| Sample | O/C (at) | C-C | $\Sigma(C-O)$ | C $sp^2$ | C-C | C-O | C=O | O-C=O | $\pi-\pi^*$ |
| --- | --- | --- | --- | --- | --- | --- | --- | --- | --- |
| GO | 0.86 | 42.4 | 57.6 | - | 42.4 | 47.9 | 3.8 | 5.9 | - |
| rGO<sub>90</sub> | 0.48 | 49.6 | 50.4 | - | 49.6 | 43.0 | 2.0 | 5.4 | - |
| rGO<sub>200</sub> | 0.10 | 76.5 | 23.5 | 75.3 | - | 18.0 |  | 5.5 | 1.2 |

In order to confirm the reproducibility of substrate preparation, we repeated AFM and XPS measurements in a representative sample of each substrate type from all batches prepared. Surface roughness was comparable in all batches tested for each particular substrate (**Figure S1**). Specifically, rGO_200_ samples turned out to be very similar intra-and inter-batches (*e.g. R_q_* values differed less than 2.5% within each particular batch and less than 17% between batches). Statistical comparisons corroborated the absence of significant differences for samples from different batches (*p* = 0.458 for *R_q_*, *p* = 0.450 for *R_a_* and *p* = 0.384 for *R_max_*) and samples from the same batch (p = 0.876 for *R_q_*, *p* = 0.791 for *R_a_* and *p* = 0.694 for *R_max_*). XPS spectra analyses also confirmed reproducibility among batches with respect to chemical composition (**Figure S2**).

### rGO_200_ prevented spontaneous differentiation of MDSCs towards pro-inflammatory phenotypes and increased their immunosuppressive capacity on stimulated splenocytes

Differentiation of spleen-isolated MDSCs (SP-MDSCs) from EAE mice towards mature myeloid cell subsets (*i.e.* macrophages, dendritic cells and neutrophils) has been previously described *in vitro* and *in vivo* resulting in their inactivation, loss of immunosuppressive capacity and EAE clinical course worsening^15^. In an attempt to use MDSCs for future cell-based therapies, and encouraged to the reported implications of GDMs on cell differentiation^22^, rGO_200_2D films were used to modulate the phenotype of SP-MDSCs *in vitro* (**Figure2A**). After 24 h in culture, rGO_200_ substrates were able to maintain SP-MDSCs in a less matured state than those cultured in conventional glass coverslips (**Figure2B-C**). Specifically, rGO_200_ substrates significantly prevented SP-MDSC differentiation towards mature macrophages (F4/80 mean fluoresce intensity-MFI-of F4/80^+^ cells over total SP-MDSCs), dendritic cells (CD11c^+^; both in percentage of cells and MFI) and antigen presenting cells (MHC-II^+^; both in percentage and MFI).

In order to determine whether maintaining the undifferentiated state of SP-MDSCs after culture on rGO_200_ could have any impact on their immunosuppressive capacity, these cells were co-cultured with myelin oligodendrocyte glycoprotein (MOG) stimulated splenocytes (1:4) obtained from EAE mice at the peak of their clinical course (**Figure2D**). As expected, SP-MDSCs cultured for 24 h on glass coverslips did not exert any immunosuppressive activity on encephalitogenic T cells (proliferation index (PI) *vs* non stimulated splenocytes, PI: MOG = 5.7 ± 0.8 and glass coverslip = 6.4 ± 0.8 %; *p* = 0.516). Remarkably, SP-MDSCs exposed to rGO_200_ for 24 h were able to significantly suppress T cell proliferation (PI: rGO_200_= 4.1 ± 0.5 %, *p* < 0.05 *vs* MOG; **Figure2E**). These data point to rGO_200_ 2D films as potent substrates to preserve the immunosuppressive capacity of SP-MDSCs by keeping them in an undifferentiated state in culture.

In an attempt to discard that rGO_200_ nanosheets could be released from the culture substrates and exert a direct effect on T cells, dynamic light scattering (DLS) analysis of the culture media was carried out. We identified a limited population of particles smaller than 100 nm in Z average (polydispersity degree-PDI = 0.2 and mean size = 10 nm). Comparatively, the hydrodynamic size of the original GO slurry in suspension was much higher (Z_average_ = 5400 nm, PDI = 0.2 and mean size = 1350 nm). These results proved that rGO_200_ nanosheets were almost absent in all culture conditions (**Figure S4**), thus ruling out major biological responses mediated by rGO nanosheets in suspension. In the same sense, previous work has shown that neither pristine nor functionalized graphene induced any direct effect on human T-cell proliferation and suppression^26^. Interestingly, a former study using an immune array of human peripheral blood mononuclear cells (PBMCs) provided convincing evidence that the adaptive immune response observed after GO exposure was not directly, but indirectly, affected by the primary effect on myeloid cells. Indeed, genes involved in direct T-cell activation, such as IL-2 and IFN-γ, were not affected, which correlated with the absence of CD69 and CD25 expression in these cells after GO exposure^27^.

### The activity state of MDSCs was substantially affected by the physico-chemical properties of 2D rGO films

We next sought to maximize the immunosuppression-enhancing effect of rGO in MDSCs by modifying the physico-chemical properties of the rGO substrates. For this purpose, we fabricated a more oxidized rGO, *i.e.* rGO_90_, by means of a milder annealing process of the GO-coated glass coverslips (90 °C for 15 min). Morphological analysis by SEM revealed comparable features to those identified in rGO_200_ substrates (**Figure S1A**). From AFM measurements, rGO_90_ 2D films characterized by *R_q_* values of 63.6 ± 7.45 nm, *R_a_* of 49.8 ± 5.60 nm and *R_max_* of 435.2 ± 55.25 nm (**Figure S1B**). Statistical analyses confirmed slight but significant differences for both *R_q_* (*p* = 0.017) and *R_a_* (*p* = 0.014) between rGO_90_ and rGO_200_, but not for *R_max_* (*p* = 0.062). More importantly, we corroborated a differential reduction degree (O/C ratio = 0.48) with respect to rGO_200_ by using XPS (**Figure S1C** and **Table 1**). Lineshape analysis of XPS spectra from rGO_90_ revealed no significant differences with that of GO slurry, where both main peaks remained, but there was a change in their relative intensity. This result is consistent with the lower O/C atomic ratio obtained and the slightly higher C-C bonding content. On the contrary, as previously outlined, C 1s emission from rGO_200_ displayed a very different lineshape dominated by an intense and asymmetric peak centered close to 284.5 eV. Taken together, these results indicate a lighter reduction of rGO_90_ than rGO_200_, supported by a significantly higher content of oxygen-containing groups in the former substrates, which remained chemically closer to the original GO slurry (Figure S3). Interestingly, just by changing the annealing temperature, we were able to prepare comparable substrates with both significantly higher surface roughness and oxygen content. It is worth to note that 2D GO films (without any thermal treatment) could not be used for cell culture due to their instability and massive sheets detachment under aqueous conditions, even at short time points of incubation.

Once characterized, the immunophenotype and immunosuppressive activity of MDSCs grown in rGO_90_ *versus* rGO_200_ were addressed. To ensure the translation of our results, we used bone marrow-derived MDSCs (BM-MDSCs) instead of SP-MDSCs for these studies. The exposition to either rGO_90_ or rGO_200_ substrates reduced both the cell percentage and MFI of CD11c^+^ and MHC-II^+^cells in BM-MDSCs (**Figure 3A-D**), similarly to results described for SP-MDSCs. Interestingly, rGO_90_ exposition dramatically diminished the percentage, not only the MFI value, of BM-MDSCs expressing the F4/80 marker, not observed before for rGO_200_. Comparing both types of rGO substrates, rGO_90_ induced a more noticeable effect than rGO_200_ on the cell percentage and MFI value for F4/80 and MHC-II markers (**Figure 3E-F**).

**Figure 3.**
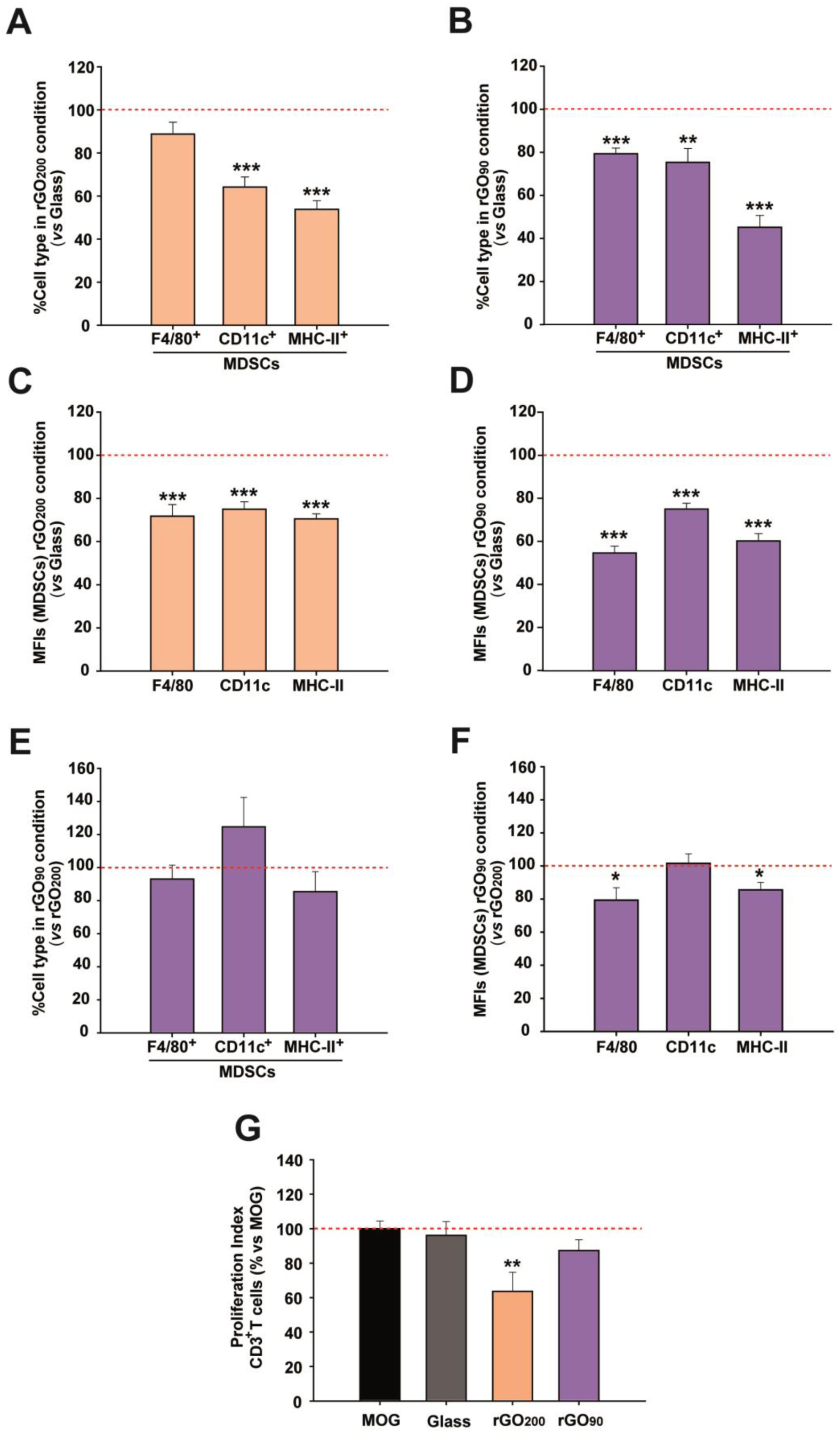
The physico-chemical properties of rGO culture substrates seem pivotal to modulate the immunosuppressive activity of BM-MDSCs. (A-D) Both rGO_200_ and rGO_90_ induced a marked reduction in the cell percentage and MFI values of cells expressing the maturation markers present in BM-MDSCs compared to glass coverslips (red dashed lines). (E-F) The effect on BM-MDSCs was more prominent when exposed to rGO_90_. (G) The exposition of BM-MDSCs to rGO_200_, but not to glass or rGO_90_, more effectively preserved their immunosuppressive activity over MOG-stimulated T cells (red dashed lines represent the proliferative activity of MOG-stimulated T cells in the absence of BM-MDSCs). Data are shown as the mean ± standard error of the mean of N =3 in immunophenotype analysis, and N = 5 for the cell proliferation analysis.

In order to investigate whether the different degree of down-regulation of specific markers induced in BM-MDSCs by rGO_200_ and rGO_90_ substrates had an impact on their immunosuppressive activity, cells were co-cultured with MOG-stimulated splenocytes from EAE mice sacrificed at the peak of the clinical course. As for the case of SP-MDSCs, isolated BM-MDSCs cultured on control coverslips lost their immunosuppressive activity over encephalitogenic CD3^+^ T cells (PI: MOG-stimulated splenocytes = 10.3 ± 1.5; glass coverslip = 7.9 ± 1.7; *p* = 0.649). Contrarily, this activity was preserved when previously exposed to rGO_200_ 2D films (PI: GO_200_ = 4.6 ± 1.5; *p*< 0.05). Remarkably, the reduction in BM-MDSC differentiation markers after rGO_90_ exposure previously described was not accompanied by a maintenance of their immunosuppressive activity (PI: GO_90_ = 11.3 ± 2.2; *p* = 0.141; **Figure 3G**). This observation is in agreement with the decreased oxidative stress and pro-inflammatory cytokine secretion observed in RAW-264.7 macrophages after exposition to suspensions of rGO nanostructures also annealed at 200 °C for 30 min^21^. In both cases, the thermal reduction of GO at 200 °C improved the biocompatibility of this nanomaterial. Therefore, our data confirm the possibility of triggering an anti-inflammatory response by GDMs, such as the site-specific Th2 response induction (*i.e.* a significant increase in IL-5, IL-3, IL-33, and their soluble receptor, sST2), as previously postulated by Wang *et al.* for graphene nanosheets intravenously administered^28^.

These data indicate that, by modulating the physico-chemical characteristics of rGO substrates, we might exert either pro-or anti-inflammatory effects on BM-MDSC activity. This finding is in agreement with previous reports on the ability of GDMs to induce changes on immune cell activity. For instance, small GO nanosheets were able to up-regulate critical genes involved in both Th1 and Th2 immune responses such as CSF2, TNF, IL6, IL10, CD80, IL1, IL1R1, TICAM1, IL8, IL23A, NFKB1, TBX21, CD40, CCR6, and IFNAR1 on human PBMCs. Also, this GO stimulated the release of some Th1 but also Th2 cytokines such as IL-1β, IL-1α, TNF-α, IL-10, IL-6, and IL-8 in the same cells^27^. In our study, although both types of rGO induced a marked reduction on MDSC differentiation markers, the more accentuated modulation after culture on rGO_90_ substrates should involve other cellular events rather than the maintenance of immaturity.

### The exposure to rGO substrates prevented the increase in cell size and complexity of BM-MDSCs in culture, being more pronounced for rGO_90_

Since alterations in the morphology of cultured peripheral myeloid cells have been associated with changes in cell functionality^29^, a morphological analysis of BM-MDSCs was next carried out in order to investigate if the differences in the immunosuppressive activity of BM-MDSCs found when cultured on rGO substrates were related to alterations in their size and shape. Consistent with the data described earlier for immunophenotyping, culture on rGO prevented BM-MDSCs from acquiring morphological characteristics of myeloid cells in the process of spontaneous maturation on glass coverslips (*i.e.* an expanded cytoplasm and presence of protrusions; **Figure 4A**). In comparison to control cells on glass, BM-MDSCs on both types of rGO films showed a significantly lower total cell as well as protrusion area (**Figure 4B-C**). In fact, exposition to rGO_200_ and rGO_90_ induced a reduction of 30.51 ± 3.36 % and 41.03± 1.71% in total cell area, together with a 60.44 ± 5.71 % and 71.04 ± 3.96 % reduction in the area of the protrusions *versus* glass, respectively. When expressed as a percentage of the total cell area, protrusions also significantly diminished in cells cultured on rGO substrates (**Figure 4D**; 38.66 ± 6.38 % and 40.72 ± 6.66 % reduction *versus* glass for rGO_200_ and rGO_90_, respectively). Similarly, the main cytoplasm area markedly decreased in comparison to glass (**Figure 4E**), showing a 12.89 ± 2.86 % and 23.37 ± 1.74 % reduction for rGO_200_ and rGO_90_, respectively. In all cases, culture on rGO_90_ induced a more pronounced shrinkage on BM-MDSCs than rGO_200_, (percentage of reduction of BM-MDSC grown on rGO_90_ *versus* rGO_200_: total cell area: 15.14 ± 2.47 %; area of the protrusions: 26.79 ± 9.99 %; percentage of protrusions area: 3.36 ± 10.86 %), although only statistically significant for the main cytoplasm area reduction in rGO_90_ *versus* rGO_200_: 12.03 ± 1.99 %). All these data indicate that rGO 2D films maintain BM-MDSCs in a less complex morphology, such effect being significantly more pronounced for the more oxidized substrate (rGO_90_). As in the case of the immunophenotype of MDSCs, the more dramatic changes induced by rGO_90_ exposure may indicate that the physico-chemical properties of these two rGO substrates (*i.e.* roughness and redox state) have a substantially different impact on BM-MDSCs.

**Figure 4.**
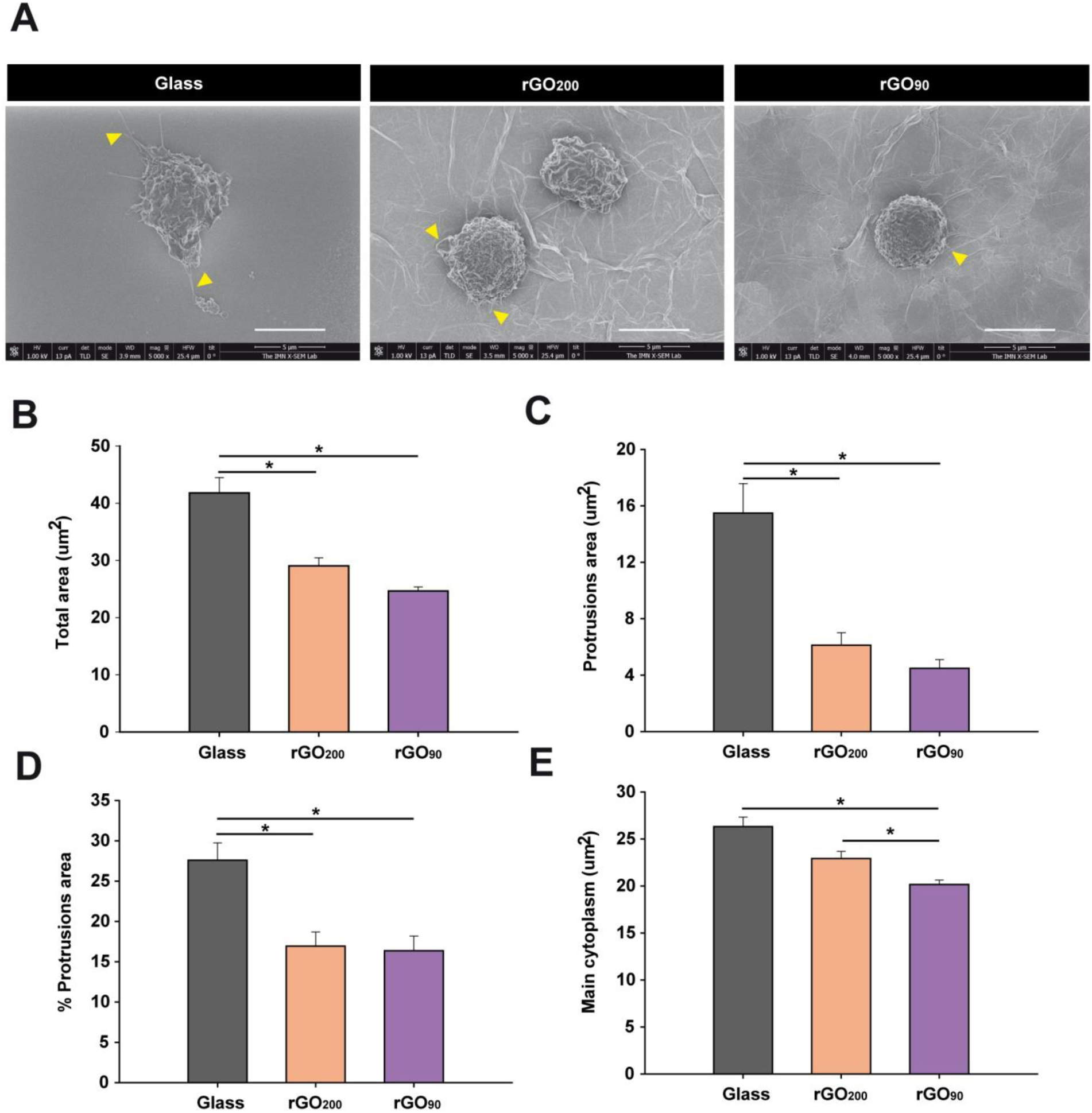
rGO substrates affect BM-MDSC size and morphology. (A) Representative SEM images of BM-MDSCs after 24 h culture on glass coverslips, rGO_200_ and rGO_90_. Arrowheads indicate examples of cell protrusions. Scale bars represent 5µm. (B-E) Both types of rGO substrates impeded cell size increase and protrusion expansion compared to cells grown on glass coverslips. Statistics: ANOVA on RANKS; p<0.001 in all cases; differences between each group versus MOG-stimulated is represented by *p<0.05 after a Dunn’s post hoc test. Data are shown as the mean ± standard error of the mean of N = 85 (glass coverslip), N = 57 (rGO_90_) and N = 66 (rGO_200_) cells from N = 2 different experiments.

In agreement with these data, Sasidharan *et al.* showed that pristine graphene impeded myeloid cells such as RAW264.7 macrophages to attain their normal stretched morphology and the formation of filopodial extensions^26^. In a different work, Hung *et al.* showed that GO significantly elicited dendritic and/or round-like morphologies in human monocytes as compared with the well-spread morphology of the control group (glass coverslips)^30^. Although our data point towards the direct physical interaction with rGO films in culture as responsible for this limitation in cell body spreading, we cannot exclude the possibility of an indirect effect of rGO. It has been previously shown that changes in myeloid cell morphology induced by GDMs are mediated in an autocrine manner^31^. When cultured in control conditioned media collected from untreated RAW264.7 cells, naïve RAW264.7 cells frequently displayed a multipolar stellate morphology, typical for macrophage polarization. However, when exposed to conditioned media collected from graphene-exposed RAW264.7 cells, naïve RAW264.7 cells often showed an apolar spherical phenotype, likely mediated by soluble graphene-induced factors. Contrarily, macrophages grown on electrochemically reduced GO presented elongated cell bodies with characteristic protrusions^32^. These controversial observations are likely related to the different physico-chemical properties of these GDMs, the cell source and phenotype, and the specific strategy followed for cell culture.

### The physico-chemical differences of rGO substrates differentially target BM-MDSC viability

Our data on BM-MDSC immunophenotyping suggest that the impairment of their morphological development seems to be related to their maintenance in a highly immature state. However, the more exacerbated reduction of phenotypic markers and smaller size presented by BM-MDSCs cultured on rGO_90_ could be also explained as a result of their entry into apoptotic pathways. In order to shed light in this hypothesis, an annexin V (AV)/propidium iodide (PrI) assay was carried out to determine whether the differential response of BM-MDSCs cultured on rGO_200_ *versus* rGO_90_substrates could be related to a modification in cell viability. First, the percentage of viable BM-MDSCs (AV^-^PrI^-^) within the total cells cultured on rGO_200_ was similar to that found on glass coverslips (**Figure 5A-B**). However, rGO_90_ dramatically reduced the percentage of viable cells after 24 h in culture. Importantly, both types of rGO substrates reduced the percentage of early apoptotic cells (AV^+^PrI^-^) compared to glass coverslips, being even lower in the case of rGO_90_ (**Figure 5C**). Finally, whereas neither rGO_200_ or rGO_90_ alter the percentage of late apoptotic cells (AV^+^PrI^+^) within the whole cultured cells, when circumscribed to the BM-MDSC population rGO_90_ promoted a remarkable increase in the proportion of dead cells compared to glass and rGO_200_ (**Figure5D**-**E**). Therefore, our results suggest that the physico-chemical properties of rGO substrates (*i.e.* roughness and, more pronouncedly, redox state) also modulate differently BM-MDSC viability.

**Figure 5.**
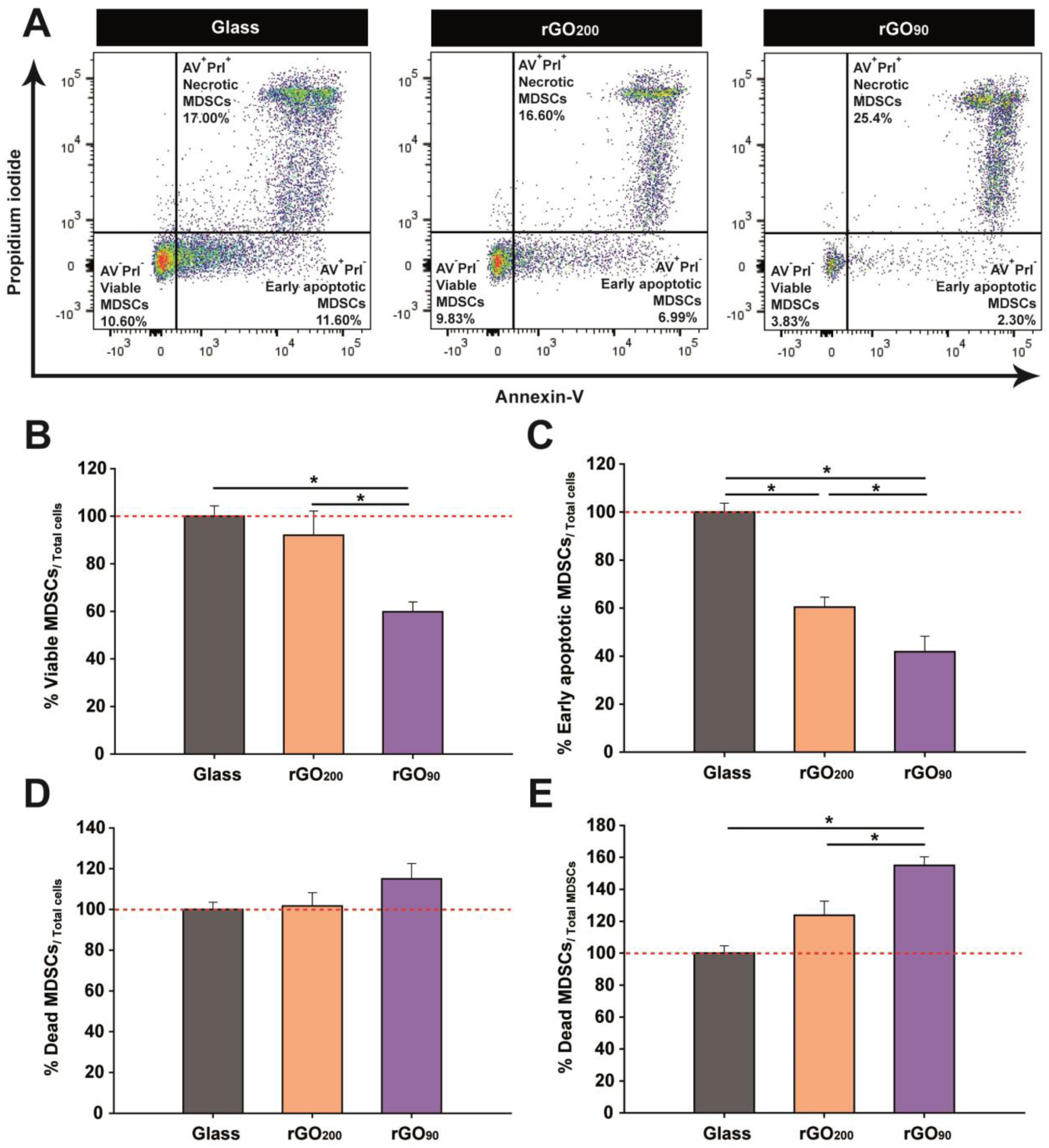
rGO substrates significantly impact viability and apoptosis of BM-MDSCs depending on their physico-chemical properties. (A) Representative dot plots of a cell viability assay with AV and PrI of BM-MDSCs cultured on glass coverslips, rGO_200_ and rGO_90_. Viable (AV^-^PrI^-^), early apoptotic (AV^+^PrI^-^) and late apoptotic/dead cells (AV^+^PrI^+^) cells were defined. (B) rGO_90_ dramatically decreased BM-MDSCs viability compared to both glass and rGO_200_ substrates. (C) rGO reduced the presence of early apoptotic BM-MDSCs, which was more remarkable in the case of rGO_90_. (D) rGO substrates had no impact on the percentage of late apoptotic BM-MDSCs with respect to total cultured cells. (E) rGO_90_ induced a statistically significant enrichment in late apoptotic cells within the population of BM-MDSCs with respect to glass and rGO_200_. Statistics: ANOVA (C-E) or ANOVA on RANKS (B) was carried out (**p < 0.01in B, ***p < 0.001 in C and E, and p = 0.121 in D); in all cases, differences between each group versus MOG-stimulated is represented by *p<0.05 after a Tukey or Dunn’s post hoc test, respectively. Data are shown as the mean ± standard error of the mean of N = 3 independent experiments.

The toxicity of GDMs on myeloid cells is still a matter of debate^33^. Some authors have reported the so-called “mask effect” to explain cell toxicity induced by GDMs^34^. Specifically, the close contact between GO nanosheets and cell membranes could drive to GO internalization. Once inside, GO could directly affect cell parameters such as viability, redox state and activation. The normal signaling and communication of cells with their environment could be also significantly altered by the presence of GO nanomaterials in their surroundings, thus influencing cell homeostasis and altering cell death, proliferation and activation cascades. In this sense, the smaller the size of GO sheets, the larger the mask effect expected. For instance, GO nanosheets (lateral dimension of 1-2 μm) in suspension were found to induce a Toll-like receptor 4 (TLR4)-mediated necrosis in different lines of macrophages, witha clear decrease on cell protrusions^35^. However, in other studies GO did not elicit any effect on cell viability and apoptosis on human monocytes after 24 h in culture. When tested *in vivo*, subdermal inoculation of GO nanosheets in a rat model increased CD163^+^ macrophages (M2-like phenotype) compared to control animals, whereas no changes in CD86^+^ macrophages (M1-like) were detected^30^. Based on these findings, and depending on GDM shape and size, myeloid cells are able to engulf and process these nanomaterials, which might either induce pro/anti-inflammatory responses or lead to the modulation and disruption of basic immune activity.

Importantly, most of the studies carried out with GDMs and immune cells to date have been performed in suspension. Indeed, the exposure to nanomaterials in solution is expected to have a larger impact on cell responses as the contact surface for interaction is maximized and they can be more easily internalized. Our particular approach deals with rGO substrates in which rGO nanosheets are integrated in stable films attached to the underlying glass coverslips. In fact, data coming from our DLS analysis proved that rGO nanosheets were almost absent in all culture conditions thus ruling out major biological responses mediated by rGO nanosheets in suspension (**Figure S4**). Indeed, several reports showed that GO 2D films could induce cell proliferation and apoptosis by cell contact, independently of cell engulfment. For instance, Escudero *et al.*, demonstrated that coating CoCr alloys with electrochemically reduced GO caused lower damage to the cell plasma membrane in J774A.1 macrophages than pristine alloys^32^. In a different study, the diminished GO internalization observed in Hri^-/-^ bone marrow-derived macrophages (with a significant deficiency in phagocytosis) did not affect cell necrosis induced by GO nanosheets^35^.

Finally, our study highlights the relevance that the physico-chemical properties of rGO play on the activation and phenotype modulation of MDSCs. In our studies, different thermal annealing conditions of GO coatings resulted in 2D rGO films with slightly different roughness and significantly different redox states. These both parameters, being the later expected to have a superior implication given its more substantial difference between rGO_90_ and rGO_200_, (Figure S3) likely participated in the modulation of MDSC responses. Besides this, concentration has been also postulated as an important factor for the control of MDSC activity^36^. Low concentrations of GO nanosheets (2.5 and 5 μg/mL) induced the proliferation of human PBMC-derived MDSCs. In contrast, a higher concentration of GO (10 μg/mL) reduced MDSC viability. In our approach, negligible quantities of rGO could be detected in suspension. Other than these properties, chemical modifications (*i.e.* functionalization) of GDMs can significantly impact their toxicity and ability to modulate myeloid cell interactions. For instance, polyethylene glycol (PEG) reduced the risk of immune responses by increasing their stability in physiological conditions and minimizing the interaction with other biomolecules. Particularly, Feito *et al.* described how PEGylated GO diminished the proliferation of macrophages in a concentration-dependent fashion^37^. PEG, in combination with polyethylenimine used to functionalize GO and served as an adjuvant, promoted dendritic cell maturation and enhanced cytokine secretion by acting over TLR-dependent pathways^38^. No data are available to date about the effect of GO functionalization over MDSCs, a field that seems pivotal for its future therapeutic use in nanomedicine.

## CONCLUSION

Our results demonstrate that 2D rGO substrates (O/C ratio of 0.10 and *R_q_* = 38.8 nm) are able to keep the immunosuppressive activity of MDSCs *in vitro*. Furthermore, by controlling the particular physico-chemical properties of GDM-based devices, we could minimize toxicity effects and take advantage of the full potential of these nanomaterials in the field of immunotherapeutics. Our findings open new perspectives on the use of rGO-based culture platforms for the expansion of MDSCs *ex vivo* as an autologous cell therapy for autoimmune pathologies such as MS. To the best of our knowledge, this is the first report of a GO-based 2D film capable of modulating the phenotype of MDSCs, a cell subset with a clear therapeutic potential. Our results also emphasize the enormous versatility of rGO nanomaterials, which are being discovered as powerful tools for biomedicine. If properly controlled, the immunomodulatory effects of rGO on MDSCs could be exploited for the treatment of diseases with a clear autoimmune etiology, such as MS. Conversely, a more oxidized form of rGO (O/C ratio of 0.48 and *R_q_* = 63.6 nm) would be an excellent nanotechnological tool in cancer, in which the elimination of MDSCs appears to be a successful strategy for the control of the exacerbated immune evasion that is behind the uncontrolled growth of tumor cells.

## MATERIAL AND METHODS

### Preparation of 2D rGOfilms

GO slurry (Graphenea, S.A.; Batch #C1250/GOB125/D; 4.6 wt% concentration, > 95 % monolayer content) was dispersed into distilled water by gentle mixing (∼10 mg/mL). This GO suspension was used to homogeneously coat circular glass coverslips (10 mm in diameter) by spin-coating at 3600 rpm for 10 s (30 µl per glass coverslip, pre-stabilization of the drop for 1 min at room temperature). The so-obtained GO 2Dfilms were then exposed to two different thermal treatments: (i) 90 °C for 15 min (more oxidized state, rGO_90_) and (ii) 200 °C for 30 min (more reduced state, rGO_200_). These treatments intend to preserve film integrity during cell culture and modify on-demand the redox state of the initial GO nanosheets.

### Physico-chemical characterization of GO slurry and 2D rGO films

TEM studies were performed by using a Jeol JEM 1010 microscope (Tokyo, Japan) at 80-100 kV with a coupled digital camera (Gatan SC200, Pleasanton, CA, USA) for image acquisition. Surface topography of rGO 2D films was visualized by using a last generation ultrahigh resolution FEI VERIOS 460 electron microscope. The analysis of the elemental and chemical composition of GO, rGO_90_ and rGO_200_ samples was performed by XPS. Experiments were carried out in an UHV chamber with a base pressure of 10^-10^ mbar equipped with a hemispherical electron energy analyzer (SPECS Phoibos 150 spectrometer) and a 2D delay-line detector, using an x-ray source of Mg-K*α* (1253.6 eV). XPS spectra were acquired at normal emission take-off angle, using an energy step of 0.50 and 0.05 eV and a pass-energy of 40 and 20 eV for survey spectra and detailed core level regions, respectively. Surface charging effect built up upon the photoemission experiments, which produces both energy shift and lineshape modifications of the spectra, has been compensated using a low energy electron flood gun. The spectra were analyzed with the CasaXPS program (Casa Software Ltd., Cheshire, UK) using a Shirley method for background subtraction and data processing for quantitative XPS analysis. Spectra are displayed after the subtraction of the contribution of the Mg-K*α* satellite emission. The absolute binding energies of the photoelectron spectra were determined by referencing to the sp^2^ transition of C 1s at 284.6 eV determined from a freshly cleaved HOPG (Highly Oriented Pyrolytic Graphite) sample. The overall sample composition was determined from survey spectra which showed intense signals from C and O, as well as a minor contributions from Na, K and S (only in GO and rGO_90_ samples). O/C atomic concentrations were determined by measuring the integral areas of O 1s and C 1s spectra, taken at a pass energy of 20 eV, after background subtraction and normalization using the sensitivity factors proportional to the Scofield cross section provided by the electron energy analyzer manufacturer^24^.

AFM studies were carried out by using a Nanoscope V forces microscope (Bruker) in tapping mode with a TESPSS tip (Bruker). Roughness values (*i.e. R_q_*, *R_a_* and *R_max_*) were obtained by using the Nanoscope Analysis software v2.0. *R_q_* is the root-mean-square (rms) value of the profile heights over the evaluation length. *R_a_* is defined as the arithmetic average of the profile heights over the evaluation length. *R_max_* is the largest of the successive values of the vertical distance between the highest and lowest points of the profile calculated over the evaluation length.

Culture media after incubation with both types of rGO culture substrates were analyzed by DLS for the detection of rGO nanosheets released from the substrates. Measurements were carried out in a Zetasizer Nano ZS instrument (Malvern). Temperature was set to 25 °C and samples were diluted with MilliQ distilled water. Data analysis was performed considering the Gaussian distributions intensity-weighted and numbered weighted, obtaining the Z average and the PDI from the first, and the mean particle size in number from the second one.

### EAE induction

Female 6 to 8 week-old C57BL/6 mice were purchased from Janvier Labs (France). Chronic progressive EAE model was induced by subcutaneous immunization with 200 µg of MOG_35-55_ (Genscript, New Jersey, USA) emulsified in complete Freund’s Adjuvant (CFA) containing 4 mg/mL of inactivated particles of *Mycobacterium tuberculosis* H37Ra (BD Biosciences, Franklin Lakes, New Jersey, USA) and at a final volume of 200 µL. Each mouse was intravenously administered 250 ng of pertussis toxin (Sigma Aldrich, St. Louis, MO, USA) diluted in 100 µL of 1X phosphate buffer saline (PBS, pH 7.4) through the tail vein the day of immunization and 2 days later. Clinical scores were evaluated on a daily basis in a double-blind manner according to the following scale: 0, no detectable signs of EAE; 1, paralyzed tail; 2, weakness or unilateral partial hindlimb paralysis; 3, complete bilateral hindlimb paralysis; 4, total paralysis of forelimbs and hindlimbs; and 5, death. Following ethical standards and regulations, humane endpoint criteria were applied when an animal reached a clinical score ≥4, when clinical score ≥3 was reached for more than 48 h or whether existed self-mutilation, persistent retention of urine, 35% weight loss and signs of stress or pain for more than 48h, even if EAE score was < 3. Animal manipulations were approved by the institutional ethical committees (*Comité Ético de Experimentación Animal del Hospital Nacional de Parapléjicos*), and all experiments were performed in compliance with the European guidelines for animal research (European Communities Council Directive 2010/63/EU, 90/219/EEC, Regulation No. 1946/2003) and the Spanish National and Regional Guidelines for Animal Experimentation (RD 53/2013 and 178/2004, Ley 32/2007 and 9/2003, and Decreto 320/2010).

### Splenic and bone marrow MDSC sorting

For SP-MDSC isolation, fresh spleens were collected from euthanized mice at the peak of their clinical symptoms defined as the day with a repeated clinical score of 3. The tissue was homogenized to a single cell suspension, passed through a 40 µm filter (BD Biosciences) and washed in Roswell Park Memorial Institute (RPMI) medium (Gibco-Thermo Fisher Scientific, Waltham, MA, USA) supplemented with 2mM L-Glutamine (Thermofisher), 10% heat-inactivated Fetal Bovine Serum (FBS,Capricorn) and 1% penicillin/streptomycin (P/S,Gibco). Erythrocytes were lysed with ACK lysis buffer (8.29 g/L NH_4_Cl, 1g/L KHCO_3_, 1mM EDTA in distilled H_2_O at pH 7.4) and the remaining splenocytes were centrifuged at 437 g for 5 min and resuspended in supplemented PBS (sPBS: 10% FBS, 25mM HEPES and 2% P/S diluted in sterile PBS).

To isolate BM-MDSCs, mice were euthanized at the peak of their clinical symptoms defined as for SP-MDSCs. Both femurs and tibiae were dissected out and the BM was flushed with supplemented RPMI media. BM cells were centrifuged at 437 g for 5 min and the obtained pellet was resuspended and red cell removed with ACK lysis buffer. After that, BM-derived cells were centrifuged at 437 g for 5 min, resuspended in sPBS, and passed through a 100 µm filter (BD Biosciences).

Splenocytes-or BM-derived cells were resuspended in sPBS and the Fc receptors were blocked for 10 min at 4°C with 10 μg/mL anti-CD16/CD32 antibodies (BD Biosciences 553142). After blocking, FITC conjugated rat anti-mouse Ly-6C (AL-21 clone), PE conjugated rat anti-mouse Ly-6G (1A8 clone) and PerCP-Cy5.5 conjugated rat anti-mouse CD11b (M1/70 clone) antibodies were added to the cell suspension and incubated for 30 min at 4°C in the dark (**Table 2**). The samples were then rinsed with sPBS, centrifuged at 371 g for 5 min, resuspended again in sPBS, and sorted in a Fluorescence Activated Cell Sorter (FACS) Aria (BD Biosciences) located at the Flow Cytometry Service of the *Hospital Nacional de Parapléjicos*. Monocytic SP-and BM-MDSCs were identified as Ly-6C^high^ Ly-6G^-/low^ gated on CD11b^+^ myeloid cells and recovered at a purity > 95%.

**Table 2.** List of antibodies used in this study for flow cytometry.

| Antibody | Target | Dilution<sup>a</sup> | Class | Clone | Manufacturer | Antibody ID |
| --- | --- | --- | --- | --- | --- | --- |
| CD11b-PerCP Cy5.5 | Myeloid cells | 0.2µg | Rat monoclonal | M1/70 | BD Biosciences | AB_394002 |
| Ly-6C-FITC | MDSCs | 0.2µg | Rat monoclonal | AL-21 | BD Biosciences | AB_394628 |
| Ly-6G-PE |  | 0.2µg | Rat monoclonal | 1A8 | BD Biosciences | AB_394208 |
| F4/80-eFluor450 | Macrophages | 0.2µg | Rat monoclonal | BM8 | eBioscience | AB_1548747 |
| CD11c-APC | Dendritic cells | 0.2µg | Hamster monoclonal | N418 | eBioscience | AB_469346 |
| MHC-II-PE-Cy7.7 | Antigen presenting cells | 0.2µg | Rat monoclonal | M5/114.15.2 | eBioscience | AB_10870792 |
| CD3e-APC | T cells | 0.2µg | Hamster monoclonal | 145-2C11 | BD Biosciences | AB_469315 |
| CD4-PE | Helper T CD4<sup>+</sup> cells | 0.1µg | Rat monoclonal | RM4-5 | BD Biosciences | AB_394585 |
| CD8a-FITC | Citotoxic T CD8<sup>+</sup> cells | 0.25µg | Rat monoclonal | 53-6.7 | BD Biosciences | AB_394568 |

<sup>a</sup>Refers to the amount per million cells. Abbreviations: FITC, fluorescein isothiocyanate; PE, phycoerythrin; PB, pacific blue.

### MDSC cultures on rGO substrates

SP-MDSCs and BM-MDSCs were resuspended in Iscove’s Modified Dulbecco’s Medium (IMDM;Biowest) supplemented with 10% FBS, 1% P/S, 50 µM β-mercaptoethanol (β-ME; Sigma), and 2mM L-Glutamine (Thermofisher). 10^5^ SP-MDSCs or BM-MDSCs were seeded on glass coverslips or rGO-based substrates. After 24h, MDSCs were harvested, centrifuged at 371 g for 5 min and resuspended in the appropriate medium for further experiments.

### Splenocyte stimulation and co-culture with MDSCs

Single-cell suspensions from the spleen were prepared as described above and splenocytes were exposed to 2-μM Tag-it Violet^TM^ Proliferation and Cell Tracking Dye (Biolegend) diluted in PBS supplemented with 0.1% bovine serum albumin (BSA) at 37°C for 20 min in darkness following manufacturer’s protocol. After washing, 2·10^5^ Tag-it Violet-labelled splenocytes were cultured in 96-well U-bottom plates (Nunclon Delta Surface; Thermo Scientific, Waltham, MA, USA) at a final volume of 200 µL of supplemented IMDM per well. Splenocytes were stimulated for 72 h with 5 μg/mL MOG_35-55_ (or culture medium alone for the unstimulated controls). To analyze the effect of MDSC growing on rGO substrates, 5 x 10^4^ MDSCs collected from glass coverslip/rGO_90_/rGO_200_ devices were exposed to pre-activated splenocytes during 24 h at a 1:4 ratio (MDSCs:splenocytes). After 48 h, all cells were harvested from the culture plates and the cell Fc receptors were blocked for 10 min at 4°C with 10 μg/mL anti-CD16/CD32 antibodies (BD Biosciences 553142). Then, cells were incubated for 20 min at 4°C in the dark with the following fluorochrome-conjugated monoclonal antibodies: anti-CD11b PerCP-Cy5.5 (M1/70 clone), anti-CD3e APC (145-2C11 clone), anti-CD4 PE (RM4-5 clone), and anti-CD8a FITC (53-6.7 clone; all from BD Bioscience). Finally, T cell proliferation, indicated by Tag-it Violet dilution, was analyzed by flow cytometry in a fluorescence activated FACS Canto II analyzer (BD Biosciences) located at the Flow Cytometry Service of the Hospital Nacional de Parapléjicos. The data obtained was analyzed using the FlowJo 10.6.2 software (Tree Star Inc.).

### Phenotypic and cell viability analysis of MDSCs grown on rGO substrates

After 24 h of culture on glass coverslips or rGO-based2D films, MDSCs were harvested, centrifuged at 371 g for 5 min, and washed in 1X PBS. Cell suspensions were then used for phenotypic and viability analyses. For myeloid markers analysis, Fc receptors from SP-or BM-MDSCs were blocked for 10 min at 4°C with 10 μg/mL anti-CD16/CD32 antibodies. Then, cells were labeled for 20 min at 4°C in the dark with the following fluorochrome-conjugated monoclonal antibodies: anti-Ly-6C FITC (AL-21 clone), anti-Ly-6G PE (1A8 clone) and anti-CD11b PerCP-Cy5.5 (M1/70 clone) all from BD Biosciences; anti-MHC-II PE-Cy7 (M5/114.15.2 clone), anti-CD11c APC (N418 clone), and anti-F4/80 eFluor450 (BM8 clone) from eBioscience-Thermo Fisher Scientific (**Table 2**). MDSCs were then rinsed with PBS, recovered by centrifugation at 371 g for 5 min, and resuspended in PBS.

For cell viability analysis, labelled BM-MDSCs were stained with the Annexin V-FITC Apoptosis Detection Kit (Beckman Coulter, IL, Italy) following manufacturer’s protocol. Briefly, BM-MDSCs were centrifuged at 93 g for 3 min, rinsed with PBS and centrifuged again to remove supernatants. Pellets were resuspended in 100 µl of binding buffer at a concentration of 10^6^ cells/mLand stained with 5 µl of AV-FITC conjugate plus 5 µl of PrIfor15 min at room temperature. The different cell staining patterns were classified as follows: 1) viable cells: double negative for AV and PrI; 2) early apoptotic cells: positive for AV and negative for PrI, and 3) late apoptotic/dead cells: double positive for AV and PrI. In all cases, MDSCs were analyzed in a FACSCanto II cytometer (BD Biosciences). The data obtained were analyzed using the FlowJo 10.6.2 software (Tree Star Inc.).

### Morphological characterization of MDSCs on 2D rGOfilms by scanning electron microscopy

Cell culture morphology was studied by SEM. Briefly, culture samples were rinsed in PBS twice and fixed with glutaraldehyde (2.5% in PBS) for 45 min. After gentle washing in distilled water, dehydration was performed by using series of ethanol solutions for 15 min (2 washes) and a final dehydration in absolute ethanol for 30 min. Samples were then dried at room temperature for at least 48 h. After mounting in stubs and coating with a nanometer-thick chromium layer under vacuum, cell culture samples were visualized by using a last generation ultrahigh resolution FEI VERIOS 460 electron microscope. From the images collected (N ≥ 50 per substrate), the impact of rGO substrates on MDSCs morphology was characterized by measuring the total cell area, the protrusions area and the main cytoplasm area using the Fiji software. To avoid biased effects, measurements were carried out by two independent observers.

### Statistical analysis

Data are expressed as the mean ± standard error of the mean. Statistical analyses were performed with SigmaPlot v11.0 (Systat Software, San Jose, CA, USA). Student’s *t*-test or Mann-Whitney U test for parametric or non-parametric data, respectively, were used to compare the effect on cell phenotype markers, T cell immunosuppression *versus* control conditions (glass or MOG-stimulated splenocytes, respectively), and rGO roughness (*R_q_*, *R_a_* and *R_max_*). An ANOVA test or its corresponding ANOVA on RANKS was performed followed by the Tukey’s or Dunn’s *posthoc* tests, respectively, to compare changes in cell morphology and viability. The minimum value of statistical significance considered was *p* < 0.05 and the results of the analysis were represented as: \**p* < 0.05; \*\**p* < 0.01; and \*\*\**p* < 0.001.

## ASSOCIATED CONTENT

### Supporting Information

The Supporting Information is available free of charge. Further details of AFM, XPS and DLS studies of different rGO substrate batches.

## AUTHOR INFORMATION

### CorrespondingAuthors

*María C. Serrano, Instituto de Ciencia de Materiales de Madrid (ICMM), Consejo Superior de Investigaciones Científicas (CSIC), Calle Sor Juana Inés de la Cruz 3, 28049-Madrid, Spain. *Diego Clemente, Hospital Nacional de Parapléjicos, Finca La Peraleda, s/n, 45071-Toledo, Spain.

### Author Contributions

CC-T: manuscript writing and most of the experimental procedures and analysis. IM-D: experimental procedures and cell quantifications. RL-G: experimental procedures related to SP-MDSCs. AGM and MCS: Preparation and physicochemical characterization of rGO substrates. FJP: XPS studies. MCS: acquisition of SEM images of MDSC cultures. DC: design of biological experiments, data evaluation and analysis. MCS and DCL: Conceptualization, manuscript writing and editing. The manuscript was written through contributions from all authors. All authors have given approval to the final version of the manuscript.

## ACKNOWLEDGMENT

The authors would like to thank Jennifer García and Inmaculada Alonso for their technical assistance, Dr Virginia Vila-del Sol and Ángela Marquina Rodríguez from the Flow Cytometry Core Facility of the *Hospital Nacional de Parapléjicos*are acknowledged for their technical assistance with flow cytometry studies. We also acknowledge the service from the MiNa Laboratory at IMN and funding from CM (project SpaceTec, S2013/ICE2822), MINECO (project CSIC13-4E-1794) and EU (FEDER, FSE) for SEM studies. FJP acknowledges financial support from Grant PID2021-126169OB-I00 funded by MCIN/AEI/ 10.13039/501100011033 and by “ERDF A way of making Europe”. The ICTS – *Centro Nacional de Microscopía Electrónica* from the *Universidad Complutense de Madrid* is acknowledged for assistance with AFM studies.

## ABBREVIATIONS

AFM, atomic force microscopy; AV, annexin V; BM-MDSCs, bone marrow-isolated myeloid derived suppressor cells; CFA, complete Freund’s adjuvant; DLS, dynamic light scattering; DMTs, disease modifying treatments; EAE, experimental autoinmune encephalomyelitis; FACS, fluorescence activated cell sorter; FBS, fetal bovine serum; GDMs, graphene-derived materials; GO, Graphene oxide; IMDM, Iscove’s modified Dulbecco’s medium; MDSCs, myeloid-derived suppressor cells; MFI, mean fluorescence intensity; MOG, myelin oligodendrocyte glycoprotein peptide; MS, multiple sclerosis; P/S, penicillin/streptomycin; PBMCs, peripheral blood mononuclear cells; PBS, phosphate buffer saline; PDI, polydispersity degree; PEG, polyethylene glycol; PI, proliferation index; PrI, propidium iodide; rGO_200_, 2D GO films thermally treated at 200 °C for 30 min; rGO_90_, 2D GO films thermally treated at 90 °C for 15 min; rms, root-mean-square; RPMI, roswell Park Memorial Institute; RR-MS, relapsing-remitting multiple sclerosis; SEM, scanning electron microscopy;sPBS, supplemented phosphate buffer saline; SP-MDSCs, spleen-isolated myeloid derived suppressor cells; TEM, transmission electron microscopy; TLR-4, toll-like receptor 4; XPS, photoelectron spectroscopy; β-ME,β-mercaptoethanol.

