## Supplementary Figures for "GRAPHENE OXIDE AS A NOVEL IMMUNOTHERAPY TOOL FOR THE MODULATION OF MYELOID-DERIVED SUPPRESSOR CELL ACTIVITY IN THE CONTEXT OF MULTIPLE SCLEROSIS"

**Figure S1.** Physico-chemical properties of rGO_90_ substrates: (A) representative SEM image, (B) AFM height image and (C) XPS C 1s spectrum.

**Figure S2.** Representative AFM height and phase images of two independent batches of rGO_90_ and rGO_200_ substrates.

**Figure S3.** Representative XPS C1s spectra from two independent batches of rGO_90_ and rGO_200_ substrates. Color code as in Figure 1 in the main text.

**Figure S4.** Hydrodynamic size distribution of rGO nanosheets released from rGO_90_ and rGO_200_ films by DLS in intensity.
